# Symmetry breaking organizes the brain’s resting state manifold

**DOI:** 10.1101/2022.01.03.474841

**Authors:** Jan Fousek, Giovanni Rabuffo, Kashyap Gudibanda, Hiba Sheheitli, Viktor Jirsa, Spase Petkoski

## Abstract

Spontaneously fluctuating brain activity patterns that emerge at rest have been linked to brain’s health and cognition. Despite detailed descriptions of the spatio-temporal brain patterns, our understanding of their generative mechanism is still incomplete. Using a combination of computational modeling and dynamical systems analysis we provide a mechanistic description of the formation of a resting state manifold via the network connectivity. We demonstrate that the symmetry breaking by the connectivity creates a characteristic flow on the manifold, which produces the major data features across scales and imaging modalities. These include spontaneous high amplitude co-activations, neuronal cascades, spectral cortical gradients, multistability and characteristic functional connectivity dynamics. When aggregated across cortical hierarchies, these match the profiles from empirical data. The understanding of the brain’s resting state manifold is fundamental for the construction of task-specific flows and manifolds used in theories of brain function such as predictive coding. In addition, it shifts the focus from the single recordings towards brain’s capacity to generate certain dynamics characteristic of health and pathology.

## 1 Introduction

The human brain at rest exhibits remarkable richness of neural activity structured both in time and space. Early computational modeling studies explored how these spontaneous fluctuations are constrained and how their organisation is shaped by the anatomic connectivity [1–4] enabling to start disentangling the mechanisms of the resting state dynamics *in silico*. A substantial body of work has related the emergent activity patterns at rest to the brain functional networks involved in task conditions [5, 6], and shown that the spatio-temporal variability of resting-state activity possesses functional significance [7–9], relevance to cognitive task performance [10], consciousness levels [11], changes during ageing [12, 13], mental disorders [14], and neurodegenerative diseases (e.g. Alzheimer’s dementia; [15]). The structure of the resting state dynamics changes over time [16] and is characterized by a range of properties such as metastability [17, 18], event-like coactivations [19–21] and traveling waves [22]. However, our understanding of the mechanisms underlying these spatiotemporal patterns of the brain activity at rest is still incomplete [23] and whole brain network models have a crucial role to play on that front [24].

There is general agreement that the resting brain operates near criticality [25]. This is supported by a large range of analyses performed on simulated and empirical data using network based measures (functional connectivity, functional connectivity dynamics), information theoretical measures (entropy, ignition) and descriptions of spatiotemporal dynamics (avalanches, cascades). Modeling efforts provide further evidence for the close relationship between the empirical data features and the properties of the structural network, local dynamics, coupling strength, neural gain [4, 13, 26–31]. The resting state dynamics can then be understood as noise-driven fluctuations of brain activity, operating near criticality and constrained by the brain connectivity [2, 32]. However, none of the above qualifies as a description of a mechanism. Descriptions of mechanisms require formulation in terms of causal activities of their constituent entities and render the end stage, in our context the resting state dynamics, intelligible by showing how it is produced [33]. To explain is thus not merely to redescribe one regularity (e.g. functional connectivity dynamics, or maximization of entropy) as a series of several (such as near-criticality, cascades, ignition). Rather, explanation involves revealing the productive relation between causal activities linked to their constituent entities.

In this paper we aim to remedy this situation and provide this explanation using Structured Flows on Manifolds (SFMs) [34–38]. SFMs is a mathematical framework explaining how low dimensional dynamics, reflecting generative sets of rules underlying behavior, emerges in high-dimensional nonlinear systems, specifically dynamical systems on networks modeling macroscale brain dynamics. When properly linked to the network’s constituent entities (functional nodes and connectivity), we will demonstrate how their causal activities lead to the formation of brain’s resting SFM, comprising all its dynamic signatures (see Figure 1). If we distill the previous reports of brain resting state data analysis from the dynamical systems point of view, we arrive at the following main empirical signatures that should be part of the end stage of a successful mechanistic description: bistability of single region activation [39–41], low-dimensionality of the global system dynamics in state space [7, 42, 43], cascade propagation [44], multistability of recurrent coactivation spatial patterns [18, 45] and their non-trivial temporal dynamics or intermittency [21, 32, 46]. These signatures will constitute the key features of what we will describe as structured flows on the low dimensional resting state manifold.

**Figure 1:**
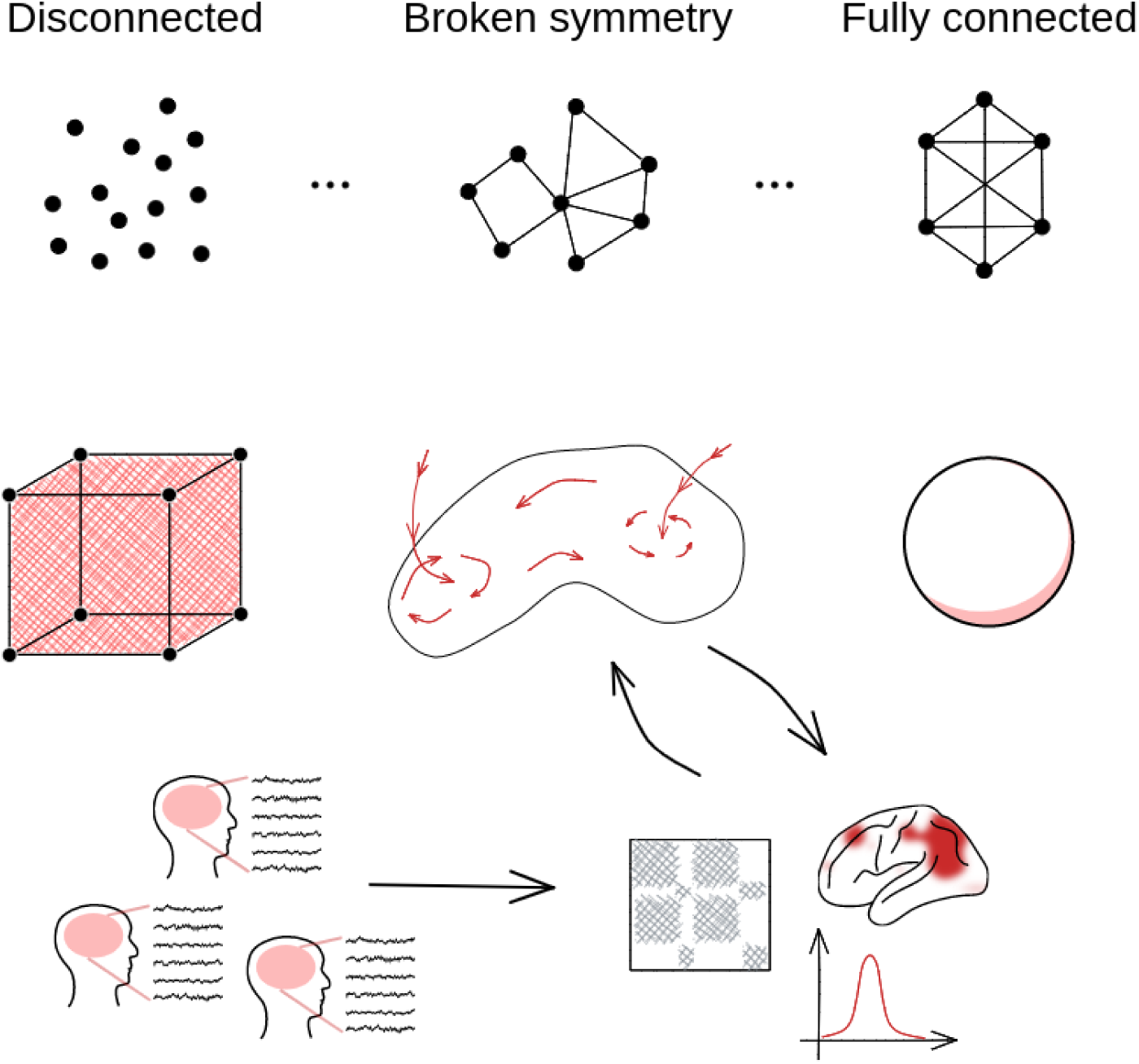
Structured flows on manifolds as focus of resting state characterization. With respect to the structure of the connectivity of the dynamical system, we consider spectrum defined by the two symmetrical limit cases: fully connected and fully disconnected network. Driven by noise, the disconnected system exhibits fully statistical, high-dimensional dynamics - it explores the whole state space in a equidirectional manner. On the other hand, the dynamics of the fully connected system is fully constrained corresponding to a SO(n) hypersphere with zero flow. The dynamics on the sparsely connected system leads to an object in between - a low-dimensional attractive manifold with an associated flow (SFM). It is this object we wish to put in the center of interest and characterize. While the SFM object remains the same, connections are made to data of various modalities with the help of suitable data features.

## 2 Results

In what follows we employ whole-brain modeling to study the low dimensional manifold and the associated structured flows of the spontaneous resting state dynamics, and how these relate to the structural connectome. We constructed a brain network model (BNM) in the Virtual Brain [47] using the two-dimensional mean-field model of an ensemble of quadratic integrate-and-fire neurons ([48]; MPR) to govern the regional dynamics coupled with a connectome derived from a subject from the Human Connectome Project [49]. We applied Balloon-Windkessel model [50] to the simulated neuronal mass activity to generate realistic BOLD signals. From these, we computed the Dynamical Functional Connectivity (dFC) that captures the changes in the system’s dynamics on the slow time scale, which we compared with empirical recordings. The fast neuronal activity is decomposed in a 2N-dimensional state space using Principal Component Analysis (PCA) to unveil the low-dimensional manifold on which the system evolves (see Materials and Methods for more details).

When driven by noise, the network of the bistable MPR nodes has the capacity to exhibit realistic dFC when the network input is scaled appropriately [44]. The noise together with the network input drives the switching between up- and down-state of the individual nodes, while the network mediates the coordination reflected in the functional connectivity. In the following sections, we explore how the manifold of the resting state activity arises from the networked interactions, how it shapes the multistability of the functional connectivity in the simulated BOLD, and how it relates to empirical observations.

### 2.1 Symmetry breaking: working point for dFC

To assess the impact of the symmetry breaking by the connectome, we simulated 10 minutes of spontaneous activity for a range of values of the coupling scaling parameter *G* and noise variance *σ*, and applied PCA to the source signal Ψ(*t*) and dFC to the BOLD (Figure 2). We used the variance accounted for (VAF) of the first two PCA components as an estimate for the dimensionality of the system’s dynamics in the state-space (Figure 2D), and the variance of the upper triangle of the dFC matrix as a measure of the *fluidity* of the system’s dynamics—that is the propensity to dwell in specific brain states (defined by the functional connectivity) and shift and return between several such states (Figure 2A). In addition, using Kolmogorov-Smirnov distance between the centered distributions of the values of the upper triangle of the *dFC_w_* in the empirical and simulated data, we have verified that the region of the parameter space where dFC is most similar to the one derived from empirical data overlaps with the region with the highest fluidity, Figure 2A.

**Figure 2:**
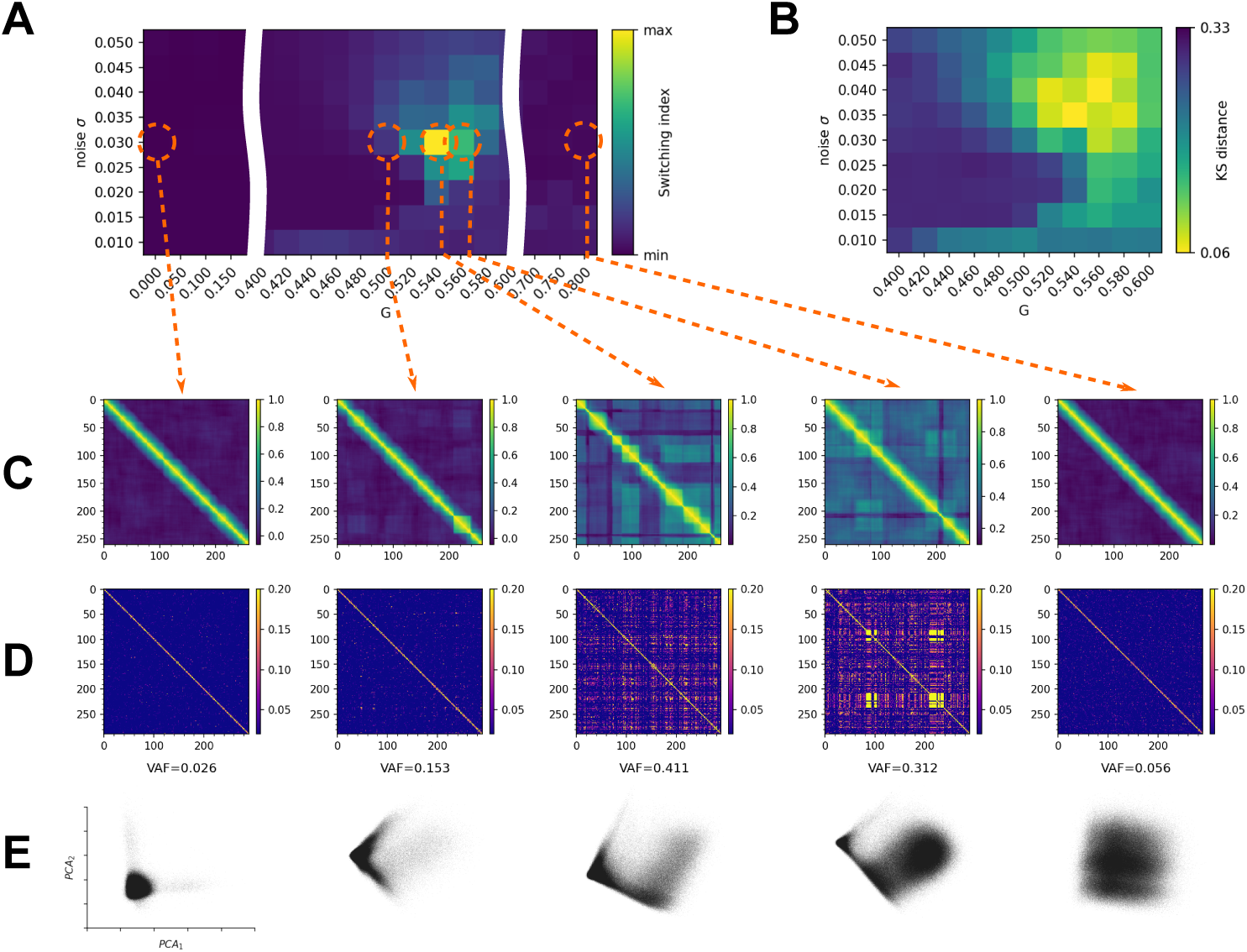
Brain network model and symmetry breaking. The brain network model is simulated for varying levels of global coupling parameter *G* and noise variance *σ* to produce both time-series of the state space variables **r**(*t*), **V**(*t*), and the BOLD signal. For each combination of *G* and *σ* we compute the sliding window *dFC_w_* matrix from the simulated BOLD signal, and quantify the ”switching index” of the dFC as the variance of the upper triangle (A). Kolmogorov-Smirnov distance of the centered (mean-subtracted) distributions of the values of the upper triangle of the dFC computed from empirical and simulated resting state BOLD time series. The region of parameter space where the distributions are closest overlaps with the region with high fluidity of dFC (B). For selected values of (*G, σ*) we show the sliding window *dFC_w_* (C), edge based *dFC_e_* (D) and the projection of **r**(*t*) time series in the first two PCA components (E) annotated with corresponding fractional variance accounted for (VAF). In the working point around *G* = 0.54 and intermediate values of *σ* the system exhibits recurrence in the large-scale dynamics as captured by non-zero switching index, and reduction of dimensionality as captured in the increase in explained variance by the first PCA components and the asymmetry in the respective projection. For values of *G* below or above the working point, the systems loses the fluidity property as reflected in the absence of the off-diagonal blocks on the dFC, and exhibits high-dimensional dynamics.

For low values of *G*, the system exhibits high-dimensional dynamics as reflected in the low variance explained by the first PCA components and low values of the variance of the off-diagonal values of dFC with the mean around 0—reflecting the absence of recurrence in the system dynamics (Figure 2B,C). Note, that the explained variance for each PCA component is equal to 1*/N* (in this case *N* = 84 nodes of the network), and the projection in panel D reflects the independent infrequent switching of two nodes, each captured by one PCA component. Around the value of *G* = *G_w_* = 0.525 and *σ_w_* = 0.030 (working point) the variance explained by the first two components of the PCA increases substantially, and so does the fluidity of the dFC as the characteristic intervals of FC invariance (on-diagonal nonzero blocks) appear together with similarity across time (high off-diagonal correlations). Past the working point (*G >* 0.6) the explained variance in PCA drops as well as the off-diagonal dFC correlations, signifying increase in dimensionality of the spontaneous dynamics.

### 2.2 Network dynamics

Before we delve into the characterization of the low-dimensional manifold, let us first describe the network dynamics in detail. For the MPR model, the dynamical profile of an isolated node in the bistable parametrization consists of an unstable fixed point (saddle node) and two stable fixed points: down-state stable node and up-state focus (Figure 3A). Considering the uncoupled system, that is, the joint dynamics of the N populations (nodes) in the absence of any inter-population synaptic coupling, the phase flow is represented by 2*^N^* stable fixed points that contain all possible combinations of the populations firing at either their low or high mean firing rates (down or up state, respectively). Starting from an initial condition, the system will settle into the nearest accessible such fixed point, a stable network state composed of a corresponding combination of regions in their up or down state. Thus, the dynamics of the uncoupled system in phase space can be thought of as being driven by a potential energy landscape with multiple stable local minima representing the stable attractor states of the network. In fact, the uncoupled system, as such, is invariant under permutation of the indexes of the populations, such that these latter attractor network states are distinguished only in terms of the respective number of nodes in up and down states. The global dynamics of the system, thus, collapses in finite time onto this stable attractor state composed of a finite set of stable equilibrium points that is invariant under shuffling of indexes of the nodes. The associated global phase flow can be decomposed into N projections onto the identical 2D phase planes of individual populations, depicted in Figure 3A. Viewed from this perspective, the structure of the basins of attraction of the 2*^N^* stable system equilibrium points redundantly inherits, in higher dimensions, the relative structure of the basins of attraction of the two stable fixed points of an individual population. For an isolated node, varying the external input *I_i_* changes the size of the basins of attraction of the stable fixed points. This modulates the probability of switching between the two states when driven by noise as captured by the mean escape times (Figure 3A, see Methods for more detail). For a connected node, the external input *I_e_* depends on the state of the neighboring nodes (see Equation 4), fluctuating as they transition between the up- and down-state. On the network level, given right scaling of the network connections, this enables the cascades of up- and down-state switching at the fast time-scale, and the co-fluctuation of the BOLD signal (Figure 3B).

**Figure 3:**
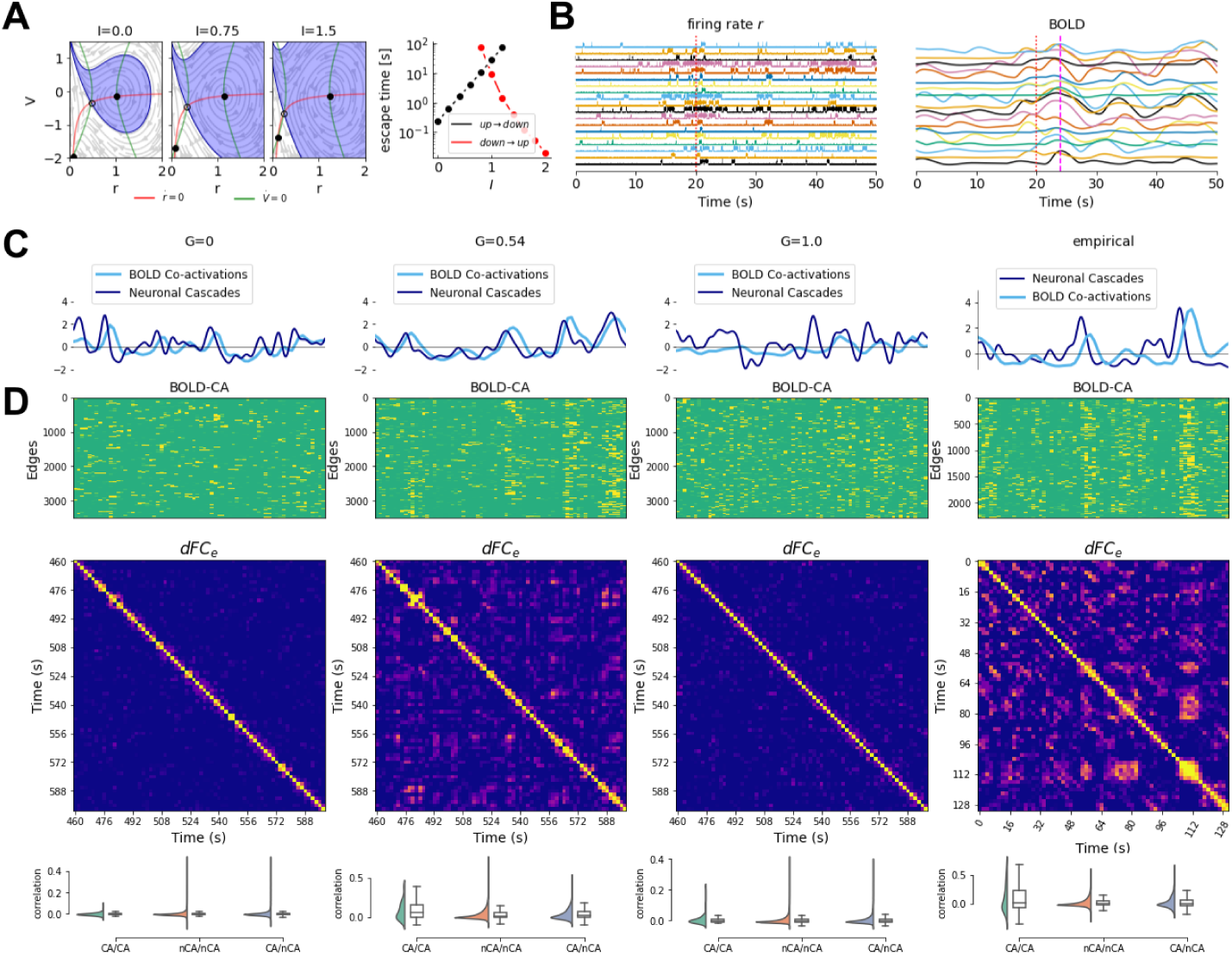
Network dynamics. (A) The network input *I* modulates the probability of a noise-driven transition between the down- and up-state. (B) Example of a cascade—coordinated increase in activity translating to a delayed correlated peak in the BOLD signal. Below we compare the network dynamics in and outside the working point, and the empirical data. In both empirical data and the working point (*G* = 0.54), the BOLD co-activations follow the neuronal cascades with a latency (C), and show distinct spatial profiles which are recurrent in time (D): edge time series on the (top panel) captures the spatial profiles of the co-activations, the similarity across time is captured by the *dFC_e_*matrix (middle panel), and the distributions of correlation between co-activation events (CA) and non-events (nCA) is compared (bottom panel).

To understand better the dynamical underpinning of the increase of fluidity of the dFC we assess the characteristics of the co-fluctuations of the BOLD signal and the cascades in the source signal. For the co-activations, we start from the edge time series which is defined as pairwise dot product of z-scored BOLD signal (an average over the edge time series would correspond to the pearson correlation). The correlation across time-points yield the *dFC_e_* matrix capturing the recurrence of the edge configurations, and the root sum squared (RSS) over the edges at each time point captures the contribution of that paricular time point to the overall functional connectivity (see Methods for more details). The time-points crossing the 95-th percentile threshold of the RSS are considered as strong co-activation events. The neuronal cascades [44] are long lasting perturbations of the neuroelectric activity and are measured on a global level as a sum over regions of the binarized firing rate activity (at the threshold of 3 standard deviations). We compared these measures between the working point *G_w_*, the disconnected system *G* = 0, the strong network coupling regime *G >> G_w_*, and the empirical data (Figure 3C,D).

In the working point *G_w_* the co-activations include large number of edges (Figure 3D) and the RSS follows the number of cascades up to a short delay corresponding to the delay of the BOLD signal. Moreover, some of the strong co-activations re-occur partially in time as reflected in the non-zero elements of the *dFC_e_* matrix. The same profiles can be observed in the empirical data, namely in the simultaneous EEG and fMRI recordings. On the other hand, the characteristic spatial and temporal structure is lost outside of the working point, that is either for the weakly coupled system (*G << G_w_*), or for too strong coupling (*G >> G_w_*).

To quantify how the co-activation events contribute to the characteristic similarity across time, we compare the correlation of the edge vectors during the events, during the non-events, and between events and non-events. As a result we observe an increased similarity of the edge vectors during the events both in the empirical data and in the simulations in the working point *G_w_*. Again, this property is lost for too weak (*G << G_w_*) or for too strong coupling (*G >> G_w_*). Together, these results show, that the system has a similar dynamical profile in the working point *G_w_* as observed in the empirical data with respect to the network-carried fluctuations on both the fast and slow timescales (as captured by *dFC* and cascades respectively).

### 2.3 Manifold of the resting state and characteristic sub-spaces

Having characterized the dynamics of the system in the working point with an appropriate measure, we proceed with the description of the manifold on which it evolves—that is to show that this behavior is low-dimensional and constrained to a specific subspace. To relate cascades and co-activations to the trajectories of the system in the 2N-dimensional state space, we first select time intervals with similar functional connectivity. Starting from the edge time series for the magnitude of co-fluctuations, we clustered the time points using k-means (k=5). This separated the high-activity intervals (majority of the nodes in the up-state), low-activity intervals (majority of the nodes in the down-state), and the co-fluctuation events (Figure 4A).

**Figure 4:**
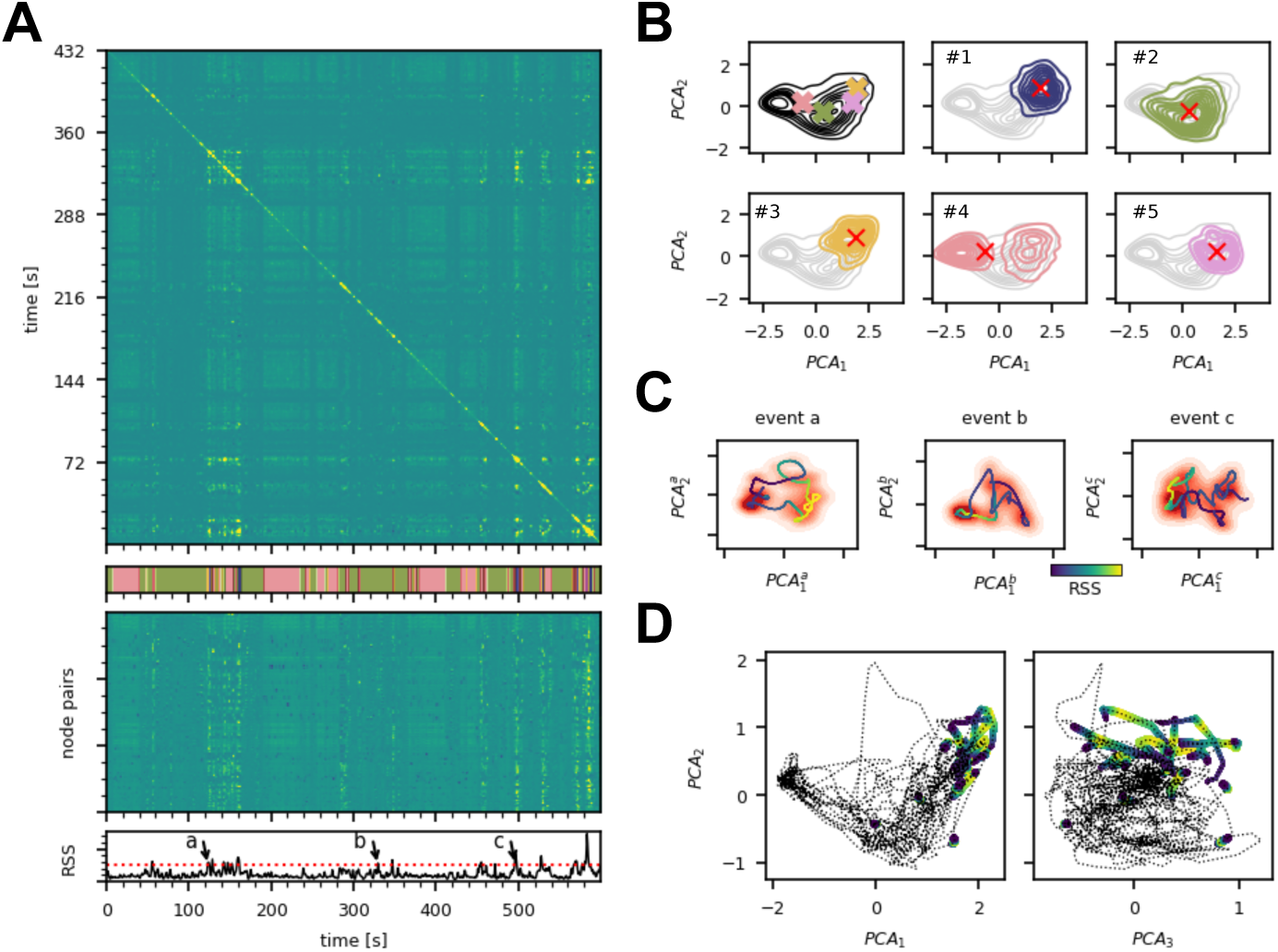
Manifold subspaces and characteristic dynamics. (A) The edge-based dynamic functional connectivity *dFC_e_* of a simulation of the model in the working point (top) shows the off-diagonal structure of similarity of the system’s activity across time. The edge time series (middle) shows the time evolution of the functional connectivity of the simulated BOLD signal between each of the node pairs, and exhibits the characteristic co-activation events defined as time points with the root sum squared (RSS, bottom) crossing the thershold of 95th percentile. Dividing the edge-time series into 5 clusters (k-means, shown in the colorbar under the dFC) has separated the event and non-event time points, and also differentiated the events based on their respective similarity. (B) The time intervals in **r**(*t*) corresponding to the 5 clusters were selected; in the first panel the centroids of the time points of the individual clusters are marked with a cross in the projection to the first two principal components of the whole time series, following panels show the projection of the **r**(*t*) intervals of particular clusters. Cluster #2 captures the high-activity subspace, cluster #4 corresponds to the low-activity state, and the clusters #1, #3, and #5 capture the co-activation events. (C) Local trajectories in the manifold subspaces: the time series of the three example events (a,b,c, marked in the panel A bottom) was projected to the first two components of PCA applied to each time segment individually. The smoothened trajectory marks the advance of the system through the event and out of it, and is colored by the value of RSS (yellow at the peak of the event blue at the beginning and the end). (D) The event trajectories on the manifold. The trajectory of the simulated BOLD signal is projected in the space defined by the first three PCA components with the events colored by the RSS value (yellow at the peak of the event).

Next, we identified the trajectories of the system underlying each cluster in the low dimensional projection of state space. For each cluster, we have selected the corresponding time points in the state space of the system, and projected them into the first two principal components of the PCA computed on the complete time series. We have observed that while the corresponding subspaces overlap partially in the projection (Figure 4B, colors correspond to the clusters), the activity within the clusters concentrates to different subspaces. This concentration in different subspaces is reflected in the distance between the centroids of the cluster time points in the PCA projection.

While the cluster activity overlaps in the projection in the components of the PCA computed from the whole time series, the co-activation trajectories become clearer by choosing different basis to span the low-dimensional space, that is, to compute the PCA from the time points corresponding to the individual co-activation events. To project the trajectories of the events observed at the slow time scale of the BOLD on the manifold, we have shifted the BOLD signal by the characteristic lag, and for each BOLD time point belonging to the cluster we selected the corresponding time points in **r**(*t*), and convolved the resulting data with a Gaussian kernel to smoothen out the noisy fluctuations (see Methods for details). We then spanned the subspace corresponding to the first two PCA components of the co-fluctuation trajectory and overlayed the smoothened trajectory over the density plot of the full **r**(*t*) time series. The density plots (shades of red, Figure 4C) of the example events show a separation of the event subspace marked by the peak in the RSS (shown in yellow in Figure 4C) on the smoothened trajectory from the rest of the manifold. This suggests that the event subspace is relatively stable, allowing the system to dwell in it long enough to cause the significant peaks in the slow BOLD signal, and that the intermediate states are less stable than the event subspace or the rest of the manifold and visited only transiently.

Although the linear embedding of the whole time-series does not separate the event trajectories well when applied to the **r**(*t*) time series, the event trajectories concentrate in the high-activity subspace spanned by the first two PCA components of the BOLD signal (Figure 4D) .

Together, these results chart the low-dimensional manifold of the system in the working point regime, associating the subspaces with specific flows. The fluid dynamics as characterized in the previous section then arise from the slow transitions between the low- and high-activity subspaces, where the latter supports the strong co-activation events which are reflected in the *dFC*.

### 2.4 Fixed point skeleton and structured flow

To understand how the resting state manifold arises, we start by considering the uncoupled system, that is, the joint dynamics of the *N* populations (nodes) in the absence of any inter-population synaptic coupling. This uncoupled system’s phase flow is dominated by 2*^N^* stable fixed points that represent all possible combinations of the populations firing at either their low or high mean firing rates (down or up state, respectively). Starting from an initial condition and in the absence of noise, the BNM will settle into the nearest accessible such fixed point, a stable network state composed of a corresponding combination of regions in their up or down state.

The dynamical effects of the symmetry breaking in the BNM are delineated by the topology of the connectome. The heterogeneity of the in-degree (total connectivity) of individual nodes of the network drives a variation in the relative positioning of the separatrices between the basins of attraction of the equilibrium points, mirrored in the variation of the corresponding projections onto the 2D phase planes of corresponding nodes (see Figure 3A). In conjunction, connectivity strength and topology give rise to gradients in the relative attractiveness of the system’s equilibrium states. This attractiveness (or stability) can be quantified by the largest negative real eigenvalues obtained from the linearization of the system about the respective equilibrium state (linear stability analysis).

To map the complete manifold outside the simulated trajectories we sampled the stable fixed points for varying coupling scaling parameter *G* from the 2*^N^*combinations of up- and down-states, and evaluated their stability (see Methods for more details). We found that the number of stable fixed points in the sample decreases with increasing *G*. This decrease is due to the loss of states with mixed composition of up- and down-state due to the bifurcation of the down state in nodes with high input (Figure 5A). Projecting the **r** component of the fixed-points in the first two eigenvectors of the Laplacian confirms this thinning of the intermediate compositions biased towards those with higher number of nodes in the up-state (corresponds to the first Laplacian eigenvector *λ*_1_). Additionally, the stability of the fixed points was inversely proportional to the number of nodes in the up-state, that is in the direction of *E*_1_ the first eigenvector of the Laplacian (Figure 5B).

**Figure 5:**
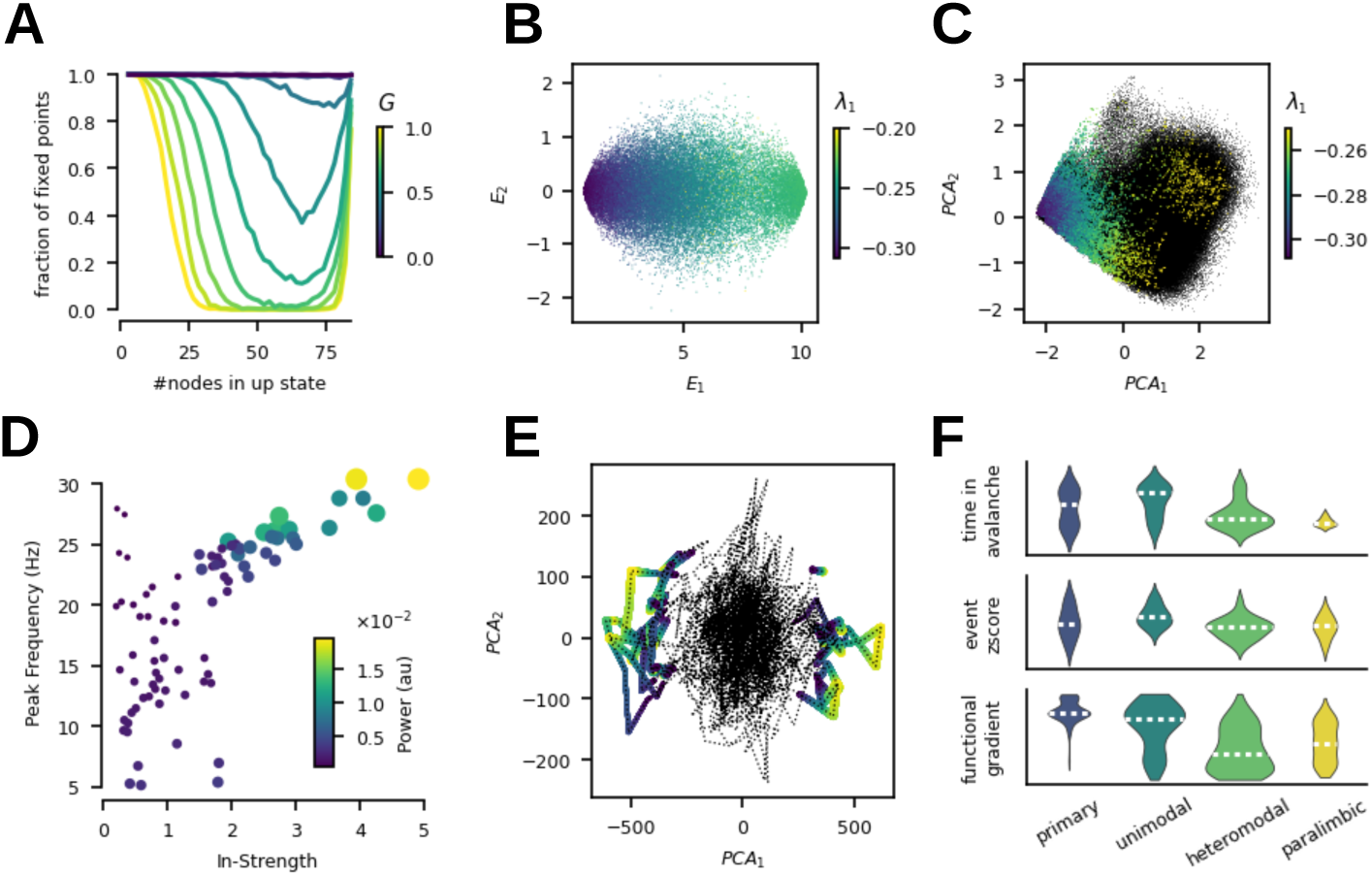
Mechanistic structure of the manifold. (A) Composition of the sampled stable fixed points in terms of number of nodes in the up-state as a function of *G*, normalized to *G* = 0. (B) Projection of the stable fixed points into the first two leading eigenmodes of the network Laplacian *E*_1_*, E*_2_, color coded with the value of the largest eigenvalue in the linear stability analysis. (B) Fixed points (colored) derived by noise-free integration to equilibrium from the trace of a simulation (black) in the working point *G_w_*, color coded by the value of the largest eigenvalue *λ*_1_. (D) Frequency peak in the simulated source activity of each of the regions plotted against the node structural connectivity in-strength. (E) Empirical BOLD time series projected into the first two PCA components with the events colored by the RSS value (yellow in the peak). (F) Across the cortical hierarchy, the time spent in avalanches of the **r**(*t*) time-series (top) decreases, as does the cumulative z-scored simulated BOLD from the event time-segments (middle). The spatial distribution of the principal functional gradient extracted from empirical fMRI is also aligned along the cortical hierarchy (bottom).

To put this into the context of the simulated trajectories, we have next identified the fixed points around which the simulated trajectory evolved by taking initial conditions from the simulated trajectory, and integrating the system without noise to the equilibrium. We have confirmed that in all instances the system reached a stable fixed point composed of combination of up- and down-states, and that the stability of these fixed points follows the same gradient in terms of the composition (Figure 5C).

Furthermore, the nodes of the network exhibit a frequency gradient of the oscillations in the up-state (Figure 5D). This gradient reflects variability of the characteristic frequency in the up-state across nodes in the network. In the fixed-point state, if the nodes are treated as isolated systems with an input current term based on the existing network state, then

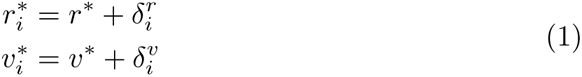

where (*r^∗^, v^∗^*) are the symmetric fixed-points of the network and 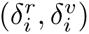 are the excursions from the symmetric fixed-point and change according to the existing network state. These excursions depend directly on the in-strength of the *i*th node and the local states of its first neighbours.

Following linear stability analysis of the *i*th system around the fixed-point (see Methods), the eigenvalues of the Jacobian matrix are given by

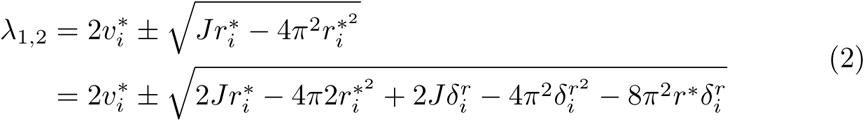

From the above equation, we see that the frequency of oscillations in the up-state of the *i*th node increases proportional to 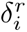 and therefore proportional to the in-strength of the node, which we also observe in the simulation (Figure 5D). Furthermore, applying the PCA projection on the empirical BOLD time-series, we have identified a similar separation of the event trajectories in the global embedding as observed in the simulations (Figure 5E). However in the case of the empirical data the system exhibits both the co-activations and co-deactivations as seen on the separation through the first PCA component.

Lastly, symmetry breaking by the connectivity alone results in a spatial organization of the above described flow which is aligned with empirically observed trends (Figure 5F). In particular, per region the time spent in avalanches and the cumulative z-scored BOLD signal within events both decrease across the cortical hierarchy from the primary to paralimbic regions. The established principal functional gradient extracted from the empirical fMRI data is also aligned along this axis .

## 3 Discussion

Using a combination of computational modeling and dynamical systems analysis we have provided a complete mechanistic description in terms of constituent entities and their causal activities leading to spontaneous co-activations and neuronal cascades in the brain’s resting state [44]. We showed how the breaking of the symmetry of the BNM’s connectivity gives rise to the structured low-dimensional dynamics in the phase space and recurrent fluctuations of the functional connectivity (Figure 2). These fluctuations arise from network-mediated cascades of up- and down-state switching and capture well the empirically found relationship between the strong co-activation events and the recurrence structure reflected by the functional connectivity dynamics (Figure 3). The subspace accessible to the brain in this regime was charted and partitioned according to the characteristic flow associated with each of the partition (Figure 4). Finally, this subspace and its associated flows arise from the rich fixed point structure of the system and the differential stability of the nodes in these fixed points is not only reflected in the propensity to state switching that reflects the cortical hierarchy, but also influences the dominant oscillation frequency (Figure 5). In summary, these results support our hypothesis that the recurrent functional connectivity states of the resting state correspond to distinct subspaces on a low-dimensional manifold associated with distinct structured flows.

The central result of our work is that the symmetry breaking via the structural connectivity carves out an attractive subspace of all the possible states of the brain, and that the flow on this manifold governs the characteristic dynamics of the brain (that is discarding the transient towards the manifold from arbitrary initial conditions). In this regime, the model captures the multistability and noise-driven exploration of the dynamic repertoire explored previously in computational studies [2, 31, 32, 51, 52]. The data features extracted from the time series provides a link between the empirical data and the model. Here, the functional structure in the brain is carried by the rare high amplitude cofluctuation events as it was previously demonstrated in empirical fMRI data [19, 21, 53], and in simultaneous EEG and fMRI measurements [44]. Similarly, recent modeling study has shown the role of structural modules of the network in shaping the co-fluctuation events [54], which is aligned with the brain network as the symmetry breaking gives rise to the low-dimensional dynamics.

The slow time scale fluctuations of the dynamical functional connectivity reflect the movement of the brain activity between the low- and high-activity subspaces of the manifold. The flow in the high-activity subspace supports the cascades, which in turn are reflected in the high activity coactivations. This movement points to the multistable rather then metastable interpretation of the resting state dynamics [55], and reflects the observation of switching between a low-amplitude incoherent and high-amplitude coherent states in empirical data [56]. Furthermore, the slow transitions between the high- and low-activity subspaces is compatible with the reports on the spontaneous infra-slow brain activity [57, 58] and the detailed reports on its spatio-temporal structure. For example, the slow traveling waves [22] propagating along the principal gradient of cortical organization [59, 60] would provide a refined description of the trajectory through the manifold subspaces.

The attractive subspace of the low-dimensional manifold and the associated structured flow arise in the presented system from the changes in the fixed-point structure due to the irregular connectivity. In particular, the network input mediates the modulation of the escape times of the noise-induced transitions. These chain into domino-like sequences [61, 62], which in turn constitute the neuronal cascades. On a network level, our results elaborate the previous analytical results of increased entropy of the attractors in an Ising-spin network model for intermediate values of coupling strength [63]. The relationship to the dimensionality of the exhibited dynamics is such that for the low values of coupling strength *G*, where the Ising model is in the trivial state with all spins equal to 0, the model presented here is also driven by noise to the all-down state due to the significantly larger basin of attraction of the down-state, and the nodes make uncoordinated noise-driven excursions to the up-state reflected in the high-dimensionality of the dynamics. For high values of *G* the situation is opposite, and for intermediate values of *G* the Ising model exhibits high entropy of attractors, which is in our case reflected in the available states organized in the low-dimensional manifold with the structured flow governed by the stability of these states.

Overall, the movement of the system through the subspaces of a low-dimensional manifold is in accordance with empirical and modeling results on recurrence and state clustering of resting state fMRI BOLD recordings. Using clustering algorithms to partition the BOLD time series yields statistically similar and temporally recurrent whole brain spatial coactivation patterns [18, 45] associated with specific dwell times and transition probabilities. However, compared to the clustering approaches applied to the BOLD time-series, the SFMs allows us to refine the partitioning of the state-space in two aspects: we unfold the subspaces based on the similarity of the coactivations on the level of the BOLD signal, and we provide a detailed description of the flow of the system in these subspaces e.g. in terms of the cascades. The former is in line with the recent advances regarding the low-dimensional representation of meso-[64, 65] and macroscopic [66, 67] brain dynamics, but the latter describes the origin of those subspaces as constrained by the connectome. Interestingly, the clustering of phase-locking BOLD states [43] leads to very similar low-dimensional representation of the resting state dynamics to our approach, with a single dominant global phase locked state and a number of transient partially phase-locked states related to functional networks. Similarly, by embedding the resting state data onto the task manifold extracted with the help of diffusion maps, [68] found that resting state time-points concentrate in the task-fixation and transition subspaces, and only a minority of time-points reach to the cognitive subspaces of the task manifold.

The description of the structured flow addresses also the fast time-scale by including the cascades, which we previously showed to relate to the co-activations observed in the BOLD signal [44]. In EEG literature, the spatio-temporal structure of the resting state dynamics is characterized with the help of microstates— sensor-level transient patterns lasting on average for 60-150 ms [69]. Attempts have been made to relate the microstates to BOLD activation clusters [70, 71], but identifying the sources generating the microstates with clustering or regression analysis has been challenging so far due to unclear relationship between the broadband EEG activity and the BOLD signal fluctuations [72]. To advance we propose to reframe the question as a search for a shared manifold of the neuronal activity with specific slow and fast time-scale characteristics which in turn are reflected in the EEG and the BOLD observables.

The manifold we describe is conceptually reminiscent of energy landscapes described in previous works [40, 41]. However, previous energy landscape models, such as in [41], implicitly assume energy minimization and thus, by construction, encode the hypothesis that the activation of two brain regions that are connected via a direct structural connection is more energetically favorable than that of two regions that are not directly connected. We make no such assumption here and, instead, the effective energy landscape emerges, in the form of a low dimensional manifold, out of the interplay of the non-linearity in the local neural mass model and the connectome, thus fully embracing the the network impact, beyond the pair-wise interactions. In addition, previous energy landscape analysis [40] assumed that the network changes only gradually by flipping one region at a time, and did not account for transitions in which several regions flip simultaneously. Treating the brain as a whole, the BNM that we presented here instead allows for such latter transitions of the system in state space, which may very well be due to strongly connected regions that are able to simultaneously influence their nearest neighbors during coactivation events.

It is worth pointing that our framework covers only one part of the mechanisms that shape the brain’s manifold and the flow on it, that is the connectome. We have assumed identical parameters for each region, ignoring the known structural hierarchies [73], which have been shown to improve the predictive value of the BNMs [56, 74, 75]. While we observed differential functional properties of the nodes across the cortical hierarchy [76], we didn’t recover the exact spatial correspondence to the established functional gradients [59]. Neuromodulation and the subcortical drives [77] are another missing aspect that similarly improve the performance of BNMs [78]. However, both of these elements are not yet established in the framework of BNMs, as is the impact of the connectome [24]. Thus our goal here is not to generate in silico observables that are as close as possible to the empirical one, which nevertheless differ a lot depending on the preprocessing, e.g. see [79], but to focus on the generative mechanisms for the key data features across time-scales and neuroimaging modalities that render functional activity identifiable across subjects [12, 13, 80].

A natural next step will be to extend the analysis to include the impact of the data-informed regional variance [81] which is now reachable by TVB through Ebrains [82]. Similarly intriguing direction for the extension of the framework presented here is in more refined inclusion of the subcortical structures, especially their impact through the neuromodulation. Notably, recent works [77, 83, 84] exploring the role of thalamus, locus coeruleus, and basal nucleus of Meynert in shaping of the dynamical landscape of the cortical activity are already formulated in the dynamical systems language while incorporating carefully the detailed anatomical and cytoarchitectural knowledge. Integrating these advances in the SFM framework is a natural next step towards the origi- nal motivation of SFM, that is to link the mesoscopic neuronal activity to the behaviour, as the intricate interactions between cortex and the subcortical areas are one of the organizing principles of the underlying the biological mechanisms supporting behaviour [85].

Parcellation-induced variation of empirical and simulated brain connectomes at group and subject levels is another issue that needs to be considered [86]. Nevertheless, we focus on general mechanisms without going on regional level specificities, so the choice of parcelation should not play such a role.

In conclusion, our results show how the low-dimensional dynamics arises from breaking the symmetry in the brain on the level of the connectome. De- scribing these dynamics as structured flows on manifolds allows us to bridge the gap between the observational measures and the state-space trajectories of the system. As such, this object is well suited for comparison across different mod- els, scales, and neuroimaging modalities, and provides means for integration of the diverse descriptions of the resting state dynamics.

## 4 Materials and methods

### 4.1 Brain network model

Computational brain network model [87] is used to simulate resting state activity under varying values of network coupling scaling parameter *G*. The dynamics of each of the network nodes were governed by the neural mass model (NMM) derived analytically as the limit of infinitely all-to-all coupled *θ*-neurons [48], namely for *i*-th node for the firing rate *r_i_* and membrane potential *v_i_* as:

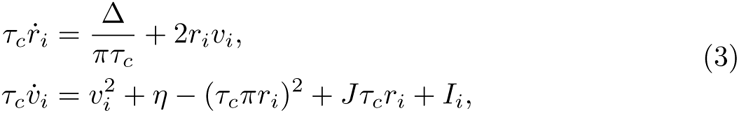

where *I_i_* is the input current, *η* is the average neuronal excitability, *J* is the synaptic weight, Δ is the spread of heterogeneous noise distribution, and *τ_c_* is the characteristic time.

The *N* nodes are then coupled with a connectome derived from empirical data as

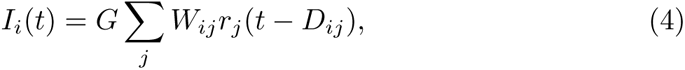

where *G* is the network scaling parameter, *W_ij_* is the connection weight, *D_ij_* = *L_ij_/S* is the delay caused by propagation of the signal on a tract of length *L_ij_* with finite speed *S*. We picked the speed *S* = 2*m/s* from the biologically plausible range [88], and a connectivity matrix of a subject from the Human Connectome Project [49] in the Desikan-Killiany parcellation [89] with 84 regions including subcortical structures.

The equations 3 and 4 comprise the drift *a*(Ψ*, t*) in the stochastic delay differential equation formulation with linear additive noise reading:

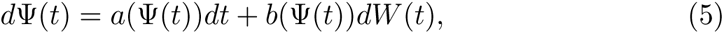

where Ψ is the state vector [*ψ*_1_*, …ψ_n_*] with *ψ_n_* = [*r_n_, V_n_*], *dW* (*t*) is a differential of a Wiener process with Gaussian increment with variance *σ*^2^, and *b*(Ψ*, t*) = 1 is the diffusion coefficient—here constant yielding the noise term additive.

The model was implemented in The Virtual Brain [47] and equipped with BOLD forward solution comprising the Balloon-Windkessel model applied to the firing rate **r**(*t*) [50].

The model parameters *η* = *−*5.0, *J* = 15.0, *τ_c_* = 1.0, and Δ = 1.0 were selected to set the nodes in the bi-stable regime in the absence of coupling [48]. We then varied the global coupling parameter *G* and the noise variance *σ*, and simulated 10 minutes of resting state BOLD activity with *TR* = 720*ms* after discarding 10 seconds of the initial transient from random initial conditions.

### 4.2 Functional connectivity dynamics

In order to track the time-dependent changes in the functional connectivity, we compute the windowed dynamical functional connectivity *dFC_w_* [32, 90] and edge dynamical functional connectivity *dFC_e_*[44]. Starting from the regional BOLD time-series *B_n_*(*t*) for each node *n*, we compute functional connectivity matrices *FC*(*w*) for each time window *w* = 1*…W* defined as 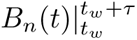 with window length *τ* = 60*s* and window step size *t*_(*w*+1)_ *−t_w_* = 2*s*. Next we compute the *dFC_w_* matrix of order *W* as

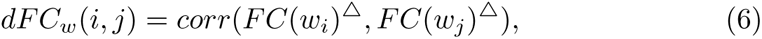

where *FC*(*w*)^Δ^ is the vectorized upper part of the *FC* matrix.

For the window-less *dFC_e_* [44] we start from the edge time-series [21] defined as 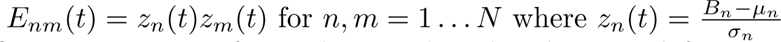 is the z-scored

BOLD time-series of a node *n*. The edge dynamical functional connectivity is then computed as correlation between the edge vectors at each pair of time points *t*_1_, *t*_2_:

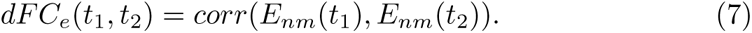

The co-fluctuation events (CF) are defined as time points in the edge time-series *E_nm_*(*t*) during which the root sum squared 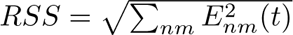 crosses a given threshold, here chosen as 95th percentile. Time points where *RSS* is below the threshold are then labeled as non-events (nCF).

The avalanches were computed on the binary mask **a**(*t*) on the **r**(*t*) such that *a_i_*(*t*) = 1 ⇐⇒ *z*(*r_i_*(*t*)) *>* 3 where *z*(*r_i_*(*t*) is the z-score of firing rate *r* of a node *i*.

### 4.3 Manifold subspaces

As a first step in the analysis of the local dynamics specific to a particular attractive subspace, we have extracted the time-points belonging to these sub-spaces with k-means clustering applied to the edge time series *E_nm_*(*t*). We varied the number of clusters *k* and selected *k* = 5 at which the co-fluctuation events separated to distinct cluster.

To extract the segments of **r**(*t*) corresponding to the *E_nm_*(*t*) time points we first estimated the BOLD signal lag *l* = 2500 ms as optimal peak-to-peak alignment with **r**(*t*) smoothened by a Gaussian filter with same effective width (*σ* = 700). Then for all BOLD time points in a given cluster *c* we selected the 2000 corresponding time points in **r**(*t*) and concatenate these to get the fast time-scale activity **r***_c_*(*t*) in the subspace corresponding to cluster *c*. Each of the **r***_c_*(*t*) was then projected to space spanned by the first two PCA components of the whole **r**(*t*) time-series to evaluate how much of the overall state-space is covered by individual clusters.

The local trajectory for a given event *e* was computed by selecting interval **r***_e_*(*t*) corresponding to BOLD timepoints above the RSS threshold and three timepoints before and after the event. Local *PCA^e^*of was then computed from **r***_e_*(*t*), and the smoothened trajectory was computed by convolving **r***_e_*(*t*) with a Gaussian filter (*σ* = 100).

### 4.4 Manifold sampling

To identify the fixed point scaffold of the manifold as traced by the trajectory resulting from integrating the Equation 5, we sample the segments from the simulated trajectory 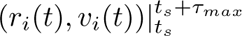, and use them as initial conditions for integration of the deterministic interpretation of Equation 5, i.e. *d*Ψ(*t*) = *a*(Ψ(*t*))*dt*. From each such an initial condition, we integrated the system to steady state equilibrium corresponding to a fixed point (**r***^∗^,* **v***^∗^*).

The number of stable fixed points (**r***^∗^,* **v***^∗^*) of system with *G* = 0 is 2*^N^* reflecting all the combinations of up- and down-states of the *N* nodes. To sample the stable fixed points of the system with *G >* 0 we solve repeatedly the system of equations:

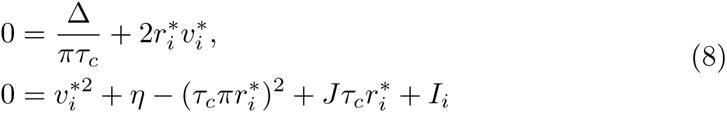

using Newton-Raphson method with the initial conditions chosen randomly as a vector of up- and down-state fixed points of the isolated nodes, i.e. 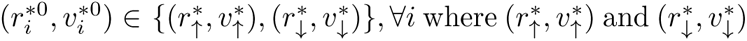 are the up- and down-state fixed points for the isolated node. For each initial condition (**r***^∗^*^0^, **v***^∗^*^0^) we then check if the corresponding solution of Equation 8 is equivalent up to the composition in terms of up- and down-states. If not, it is discarded, otherwise we evaluate the stability of the found fixed point using linear stability analysis.

As a low-dimensional projection of the sampled manifold we have used the two slowest eigenmodes of the structural connectivity. These are computed as eigendecomposition of the graph Laplacian **L** = **W***−***I**, that is **LU** = **UΛ**, where eigenvalues *λ_k_* can be interpreted as structural freqencies and the eigenmodes

**u***_k_* as structural connectome harmonics [91].

### 4.5 Linear stability analysis

We perform a linear stability analysis to identify the fixed-points obtained from the NR method. If each fixed-point (**r***^∗^,* **v***^∗^*) is perturbed by (ɛ^r^, ɛ^v^), then the evolution of the perturbations depend on the Jacobian matrix (*J*) and are given by:

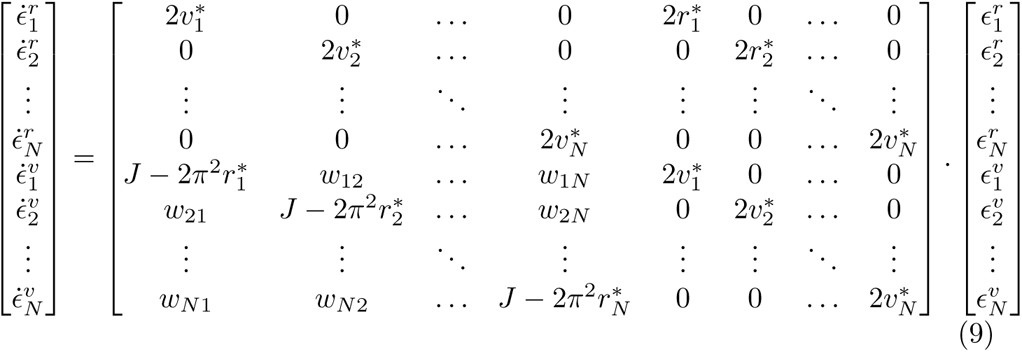

The stability of a fixed-point depends on the eigenvalues of the Jacobian evaluated at the fixed-point. The fixed-point is stable iff all the eigen-values of *J* are negative. Therefore, we numerically evaluate the largest eigenvalue of Jacobian for each fixed-point and label the point as stable if its real-part is negative.

### 4.6 Fixed point sampling from simulated trajectory

From a given trajectory of the system given as 10 minutes of *ψ*(*t*) we have selected a restart point *t’* each 50 ms (12000 starting points altogether). For each of the restart point *t’* we extracted the segment 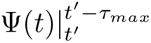 where *τ_max_* is the length of the longest delay, and used as initial condition to a equivalent system to Equation 5 with *b* = 0:

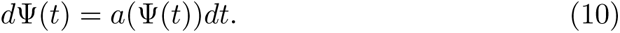

Integrating this system to equilibrium yielded then for each restart point *t_r_* a fixed point Ψ*^∗^* = (**r***^∗^,* **v***^∗^*). The stability of each of the fixed points Ψ*^∗^* was then evaluated using the linear stability analysis as the largest eigenvalue of the respective Jacobian matrix.

### 4.7 Escape time analysis

The switching behaviour of a single node is driven by the stability of the up- and down-state fixed points in the presence of noise. We employed escape time analysis [92] to measure the stability of these fixed points for range of values of external input *I*. In detail, for a single node of the system given by Equation 3 we found the up- and down-state stable fixed points (*r^∗^, v^∗^*)*^↑^* and (*r^∗^, v^∗^*)*^↓^*, and the unstable saddle node (*r^∗^, v^∗^*)*^×^*. Next we computed the separatrix between the two basins of attraction by integration of the model backwards in time resulting in an closed curve *ω*. To find the characteristic escape time for a fixed point (*r^∗^, v^∗^*) we have integrated the system from the initial condition (*r*_0_*, v*_0_) = (*r^∗^, v^∗^*) for a given value of *I* 100 times, measuring the time *t_E_* at which the trajectory crosses *ω* for the first time. The values of *I* were drawn from the range given by [0*, I*_max_] where *I*_max_ = max{*I_i_*(*t*)*, ∀i*} is the largest value of *I_i_* encountered in the integration of the full system in the working point.

### 4.8 Empirical data and spatial analysis

The functional gradient on empirical data was computed from the group connectivity matrix of the Human Connectome Project dataset using the brainspace toolbox [93]. For a simulated resting state session with *G_w_*, the time in avalanche was computed for each node as total time for which the **r***_i_*(*t*) was above the threshold of 3 standard deviations, and the event z-score as a sum of z-scored BOLD signal in time-points marked as events. The nodes were then groupped according to the cortical hieararchy [76] projected to the Desikan-Killiany parcellation.

A parcellation-based BOLD signals of a resting-state session from a subject from the Human Connectome Project [94] were used to validate the separation of the events in the low-dimensional embedding. The data consisted of 1200 time points sampled at 720 ms in the Desikan-Killiany parcellation [89] with 70 cortical regions.

